# X-ray structural analysis of single adult cardiomyocytes: tomographic imaging and micro-diffraction

**DOI:** 10.1101/2020.07.08.193771

**Authors:** M. Reichardt, C. Neuhaus, J-D. Nicolas, M. Bernhardt, K. Toischer, T. Salditt

## Abstract

We present a multi-scale imaging approach to characterize the structure of isolated adult murine cardiomyocytes based on a combination of full-field three-dimensional (3d) coherent x-ray imaging and scanning x-ray diffraction. Using these modalities, we probe the structure from the molecular to the cellular scale. Holographic projection images on freeze-dried cells have been recorded using highly coherent and divergent x-ray waveguide radiation. Phase retrieval and tomographic reconstruction then yield the 3d electron density distribution with a voxel size below 50 nm. In the reconstruction volume, myofibrils, sarcomeric organisation and mitochondria can be visualized and quantified within a single cell without sectioning. Next, we use micro-focusing optics by compound refractive lenses to probe the diffraction signal of the acto-myosin lattice. Comparison between recordings of chemically fixed and untreated, living cells indicate that the characteristic lattice distances shrink by approximately 10% upon fixation.

**SIGNIFICANCE:** Diffraction with synchrotron radiation has played an important role to decipher the molecular structure underlying force generation in muscle. In this work, the diffraction signal of the actomyosin contractile unit has for the first time been recorded from living cardiomyocytes, bringing muscle diffraction to the scale of single cells. In addition to scanning diffraction, we use coherent optics at the same synchrotron endstation to perform holographic imaging and tomography on a single cardiomyocyte. By this hard x-ray microscopy modality, we extend the length scales covered by scanning diffraction and reconstruct the electron density of an entire freeze-dried cardiomyocyte, visualizing the 3d architecture of myofibrils, sarcomers, and mitochondria with a voxel size below 50 nm.

## INTRODUCTION

Force generation in heart muscle relies on a hierarchically ordered structural organization, reaching from the actomyosin assembly in the sarcomere to the entire structural organization of cardiomyocytes (CMs). The latter includes, for example, the dense packing of myofibrils, the high number of mitochondria, and the structural and dynamical properties underlying excitability. Classical x-ray diffraction studies of skeletal (1) and heart muscle (2) have helped to shape our understanding of the average structure of the sarcomere. In contrast to electron microscopy, muscle structure analysis by x-ray diffraction is compatible with *in situ* mechanical loading, physiological parameters, and simultaneous measurements of the contractile force. Many molecular details of the myosin head dynamics, binding, and stroke have been revealed by third generation synchrotron radiation (3–6). In these experiments, however, structural information is averaged over macroscopically large volumes of the muscle tissue, without sensitivity to the cellular organization, or cell-to-cell variations. Electron microscopy, on the other hand can unravel the molecular and sub-cellular organization of myocytes (7, 8), but requires invasive staining and sectioning, while confocal fluorescence microscopy is compatible with *in vivo* recordings of contracting cells (9), but lacks the resolution for the molecular scale and also contrast for unlabeled structure.

With recent progress in x-ray focusing optics, as reviewed in (10), it has become possible to perform diffraction experiments with spot sizes in the micron and nanometer range. This enables recordings of the small-angle x-ray scattering (SAXS) while scanning the sample in real space, as initially introduced for biomaterials (11, 12), more recently also for soft matter (13), as well as for 3d vector SAXS (14). Scanning diffraction from biological cells with spot sizes smaller than a single organelle are still more challenging in terms of signal-to-noise (15–20), as well as radiation damage (21). Recently, we have used this approach to study the cytoskeletal structure of single CMs in order to compare the signal level for different preparation states (22). For freeze-dried heart muscle cells (20, 23), we have observed anisotropic diffraction patterns, and by correlating the x-ray signal to fluorescence data obtained by *in situ* stimulated emission depletion (STED) microscopy recordings, we have linked the signal to the actin portion of the cell (24). A diffraction signature of a sarcomeric complex was observed for the first time for hydrated and chemically fixed cardiomyocytes (22). Hence, by this technique, variations of structural parameters in single isolated cells or in muscle tissue have now become accessible, unaffected by macroscopic averaging. However, it cannot be expected that the signal level and amount of structural details can compete with macroscopic muscle diffraction.

Regarding sub-cellular and cellular organization, a second line of development, not in x-ray diffraction but in coherent imaging with hard x-rays has also matured (25–28), providing 3d electron density maps for unstained cells, reconstructed either by coherent diffractive imaging (29), ptychography (19, 30, 31), or by x-ray holography (32–35). While the resolution is still lower than in x-ray microscopy with Fresnel zone plate lenses (36), the size of adult CMs is too large to be penetrated by soft x-rays in the water window.

In this work, we now combine full-field 3d imaging of the electron density by holographic x-ray tomography (holotomography) and scanning x-ray diffraction to study the structure of isolated murine adult CMs on the molecular and cellular scale. As illustrated in Fig. 1, this multi-modal x-ray approach has been implemented at the same synchrotron endstation, where both modalities can be realized by a simple rearrangement of its modular compound optics (37). The purpose of the current work is threefold. First, this study is a benchmark of current x-ray method development geared towards cellular biophysics and cell biology. Second, it can contribute to computational modeling of the contractile functions of CMs. Third, it illustrates the capability and limitations how x-ray imaging methods can contribute to future multi-modal correlative imaging approaches. Such efforts will most certainly comprise visible light, electron microscopy and fluorescence microscopy, and hopefully also cover structural dynamics. To this end, the ‘x-ray contrast’ offering quantitative electron density as well as actomyosin lattice parameters without slicing or staining, will be complementary to the other more established microscopy methods. The following achievements highlight the progress that has been made with respect to prior work: (i) The first demonstration of diffraction analysis for (initially) living cells, which shows a proper exploitable acto-myosin signal. (ii) The demonstration how the actomyosin lattice spacing changes when the CMs are chemically fixed. (iii) The first 3d electron density reconstruction by phase contrast x-ray tomography, albeit at this time only for a freeze-dried cell.

**Figure 1:**
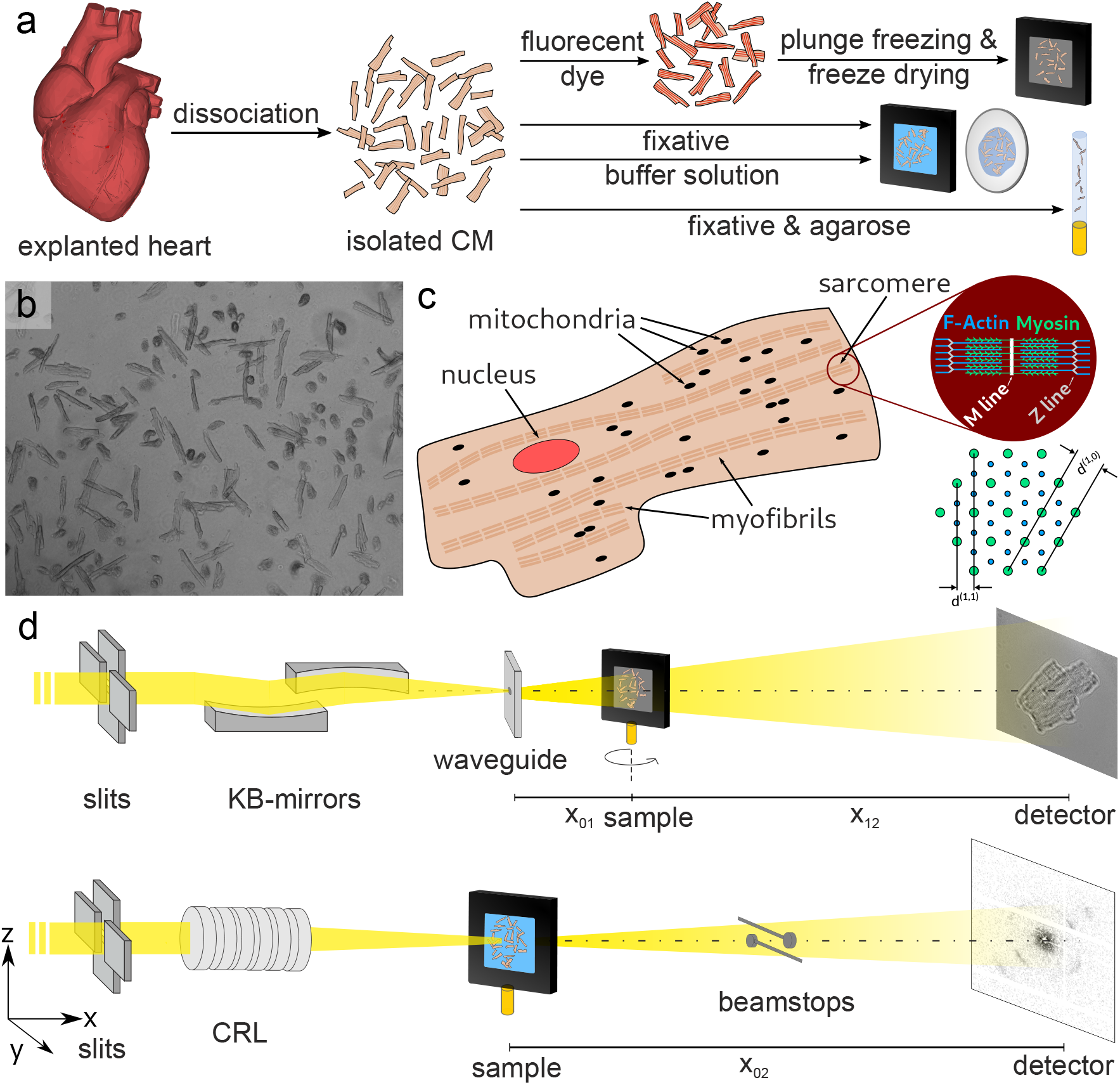
(a) Sample preparation. CMs were isolated by dissociation of healthy mouse hearts. One part (top row) of the cells was chemically fixed, mounted on a silicon nitride window and actin was stained with a fluorescent dye (Atto633-Ph). Afterwards, the samples were plunge frozen and freeze dried. For experiments in hydrated conditions (middle rows), the cells were either analyzed in buffer solution or chemically fixed. They were placed in liquid chambers made of SiN (chemically fixed cells) or polypropylene windows (living cells). For tomography, fixed cells were mixed and embedded in agarose and mounted in a polyimide tube with a diameter of 1 mm (bottom row). (b) Microscopy image of freshly isolated CMs. Healthy, contracting cells and apoptotic cells can be identified. (c) Schematic overview of the cellular structures of a CM. The nucleus, mitochondria and myofibrils are sketched. The red circle marks a sarcomeric unit which is magnified in the scheme on the right. A cross section of the sarcomere and the definition of the sarcomeric lattice spacing is shown below. (d) Experimental setup for x-ray phase contrast tomography at the GINIX endstation of the P10 beamline at the PETRAIII storage ring (DESY, Hamburg, Germany). The x-ray beam is focused by a pair of Kirkpatrick-Baez (KB)-mirrors and coupled into a waveguide channel for further reduction of the beam size and for coherence filtering. The sample is placed on a fully motorized tomography stage at distance *x*_01_ ≈ 0.35 m and the detector is placed at *x*_12_ ≈ 5 m behind the sample. The resulting cone beam geometry of the highly coherent wavefront is used to acquire projections at high geometric magnification *M* ≫ 1 and low Fresnel number *F* ≪ 1. (e) The same endstation can also be used for scanning diffraction experiments. In this mode, the beam is focused by CRLs and the sample is moved into the focus. Diffraction patterns are acquired by an EIGER 4M detector at distance *x*_02_ ≈ 5 m. The primary beam is blocked by beam stops to protect the detector.

The manuscript is organized as follows. Following this introduction, we describe methods of sample preparation, instrumentation, optics, reconstruction, and analysis. We then present the holo-tomography results for single CMs, followed by the x-ray micro-diffraction results for living CMs, before closing with a brief summary and outlook.

## MATERIALS AND METHODS

### Preparation of CMs for x-ray analysis

The workflow of the sample preparation is sketched in Fig. 1a. After the dissociation of the hearts, single CMs were prepared for the x-ray analysis. The structure of CMs, as sketched in Fig. 1c, was analyzed using the two modalities of the beam line setup. For the investigation of sub-cellular structures such as the nucleus and mitochondria, full-field holographic x-ray tomography was used, while the molecular actomyosin structure (sketched in the magnification of Fig. 1c) was probed by scanning x-ray diffraction. For diffraction measurements, the CMs were mounted in liquid chambers, whereas for tomographic scans the cells were either freeze dried or embedded in agararose to increase stability. A detailed description of the preparation steps is given below.

### Isolation of cardiomyocytes

The isolation of adult CMs followed the procedure described in (38). Before sacrifice, the wild type mice (C57BL/6) were anesthetized with isoflurane. The hearts were explanted and put in an ice-cold tyrode-solution to wash out remaining blood. After this, a canula was put in the aorta and the heart was washed with the tyrode solution. The hearts with the canula was put into a Langendorff perfusion system containing a liberase solution which was pre-heated to 36.5° C. The hearts were perfused by this solution, starting the digestion of the heart tissue. After 5 minutes, the hearts were transferred to a Petri dish, also containing the enzyme-solution. The hearts were cut into smaller pieces and resuspended in the same medium using a 10 ml pipette with a cut off tip. The resuspended cells were mixed with a stop solution (containing bovine calf serum (BCS) and CaCl_2_) in a Falcon tube. The remaining tissue sinks to the bottom, the supernatant is transferred to another falcon tube and the cells sediment and form pellets. Afterwards, the cells were washed twice with tyrode-solution. Directly after the isolation, the extracted cells were observed in the microscope. The success of the isolation was evaluated by the vitality of the cells. Figure 1b shows a microscopy image of freshly extracted CMs. The healthy, contracting cells can be distinguished from apoptotic cells. Only isolations with a vitality rate above 50% were used for further analysis.

### Preparation of CMs for diffraction

For the diffraction experiments, the isolated CMs were transferred to liquid sample chambers, see middle row of Fig. 1a. Part of the cells were chemically fixed right after the extraction. Karlsson-Schultz (KS) fixation solution (39) containing 4% formaldehyde and 2.5% glutaraldehyde was used as a fixative. The cells were incubated in the solution for 15 minutes and stored in PBS at 4° C for 2 weeks before the experiment started. For the x-ray diffraction measurement a chamber from Silson Ltd. (Warwickshire, United Kingdom) was used. The chamber consists of two 1 *μ*m thick silicon nitride membranes in silicon frames. One of the frame has a 70 *μ*m polymer spacer, keeping the frames apart in order protect the CMs from squeezing. The cells, with a diameter of approximately 10-25 *μ*m and a length of 50-100 *μ*m, are carefully sandwiched between the two silicon nitride frames. To seal the chamber, the silicon nitride frames are placed in a metal frame, which is sealed with two gaskets and screwed together. The metal frame is then inserted into a sample holding pin. The investigation of living CMs required a higher effort in the planning of the experiments. Right after the extraction, the living CMs were transported from Göttingen to the beamline in a mobile incubator (37°C). The cells were stored in tyrode buffer during transport and measurement. At the beamline the living cells were sandwiched between two polypropylene membranes. To keep the membranes apart, a gasket of the silson chambers with a thickness of 300 *μ*m were used. The polypropylene membranes were sealed tightly by applying nailpolish to the edges. The chamber was then inserted into a sample holding pin for the measurement. Additionally, plunge frozen and freeze-dried CMs mounted on SiN windows were analyzed. Since this preparation was primarily used for holographic and tomographic imaging, the procedure is described in the next paragraph.

### Preparation of CMs for x-ray holography and tomography

For the tomographic scans, CMs were either coated to SiN windows, plunge frozen and freeze dried (top row of Fig. 1a) or embedded and hydrated in 1% agarose gel (bottom row of Fig. 1a) to increase stability. In order to reduce absorption for the agarose embedded cells and to allow for a rotation of the sample, the cells were transferred into a capillary using a 1 mm biopsy punch. The protocol of the preparation for freeze-dried samples closely followed (20, 22, 23). CMs were pipetted onto SiN-windows coated with a 1% fibronectin coating and incubated for 15 min. Afterwards the cells were chemically fixed (15 min in 7% formaldehyde). Additionally, the actin skeleton of the CMs was stained with a fluorescent dye (Atto633-phalloidin, ATTO-TEC GmbH, Siegen, Germany). Plunge freezing of CMs was performed with a grid plunger (Leica EM GP). Afterwards, the cells were freeze dried. The stained samples were stored in a light-protected desiccator until the measurements were performed. As an alternative to freeze-dried preparations which compromise the structural preservation of the ultra-structure, we also used embedding in a gel to prevent dehydration, while at the same time increasing positional stability for the tomographic scan. Since the native electron density contrast in hydrated CMs was found insufficient, contrast enhancement by staining, namely by 1% OsO_4_ and phosphotungstic acid (PTA) (40, 41) was used.

### Inspection of CMs by STED

Before x-ray holography and tomography experiments were performed, the freeze-dried CMs were imaged with a custom build STED microscope (Abberior Instruments, Göttingen, Germany)(24). This setup is compatible with the GINIX endstation of the P10 beamline. For the beamtime block of the current work, however, the STED microscope was only available in ‘offline mode’ at the institute for x-ray physics in Göttingen. It allowed to pre-characterize the freeze-dried samples before the beamtime. In this way, cells which did not show any ruptures from the procedure of plunge freezing and freeze drying were selected for x-ray imaging. First, the cells were analyzed in confocal mode and raster scanned with a step size of 200 nm in lateral and 400 nm in axial direction. Additionally, one slice was acquired in STED mode and a step size of 50 nm.

### Synchrotron beamline and instrumentation

The x-ray experiments were carried out at the undulator beamline P10 at Deutsches Elektronen-Synchrotron (DESY) in Hamburg, Germany, using the GINIX (*Göttingen instrument for nanoscale imaging with x-rays*) endstation (37). The modular design of the GINIX endstation enables rapid switching between the two modalities used here: holo-tomography based on a Kirkpatrick-Baez (KB) and x-ray waveguide compound optics (32, 42, 43) and scanning diffraction based on micro-focusing by a Beryllium compound refractive lens (CRL) transfocator system. Both settings are sketched in Fig. 1d. Experimental parameters, concerning the instrumentation and optics, are tabulated in Tab. 1 and described in more detail below.

**Table 1:** Beamline specifications of the setups for scanning SAXS and tomography of the GINIX endstation.

|  | sSAXS | tomography |
| --- | --- | --- |
| energy (keV) | 8 | 7.5 |
| monochromator | Si(111) channel-cut | Si(111) channel-cut |
| detector | Eiger 4M | sCMOS (15 $\mu\text{m}$ Gadox scintillator) |
| source-to-detector distance $x_{02}$ (cm) | 507.5 | 501.14 |
| pixel size $px$ ($\mu\text{m}$) | 75 | 6.5 |
| focus (horz. × vert.) | $2 \times 3 \mu\text{m}^2$ | $< 50 \text{ nm}$ |
| $I_0$ (photons/s) | $2.5 \cdot 10^{10}$ | $1.1 \cdot 10^9$ |

For x-ray diffraction, the undulator beam was monochromatized by a Si(111) channel-cut crystal to an energy of *E*_ph_ = 8 keV. The beam was focused by a set of Beryllium CRLs to a size of 2 × 3 *μ*m^2^. The total photon flux was 2.5 · 10^10^ photons/s. The sample was placed in the focal plane and moved by a fully motorized sample stage using a piezo-motor. To protect the detector, the primary beam was blocked using two beamstops. The diffraction signal was recorded by a single photon counting detector with a pixel size of *px* = 75 *μ*m (Eiger 4M, Dectris, Switzerland) located approximately 5 m behind the sample.

For the holographic and tomographic scans, the photon energy was set to *E*_ph_ = 8 keV and 7.5 keV in two subsequent beamtimes. The beam was focused by a set of Kirkpatrick-Baez (KB) mirrors to a focal spot size with a full width at half maximum of about FWHM_y,z_ = 300 × 300 nm^2^ in horizontal (*y*) and vertical (*z*) direction. The photon flux was *I*_0_ ≈ 1.1 · 10^11^ photons/s. The KB-focused beam was then coupled into a lithographic x-ray waveguide channel (44, 45), which is placed in the focal plane. The waveguide serves the purposes of further beam confinement as well as spatial and coherence filtering, providing a clean wavefront for inline holography (32). At the detector plane a photon flux of *I*_0_ ≈ 1.1 · 10^9^ photons/s was detected. The sample was moved by a fully-motorized sample stage to a defocus plane and probed by the highly coherent exit wave field emanating from the waveguide. Small phase shifts of the wavefront after interaction with the object transform into a measurable intensity pattern by free space propagation of the beam and self-interference of the divergent wave field. The geometrically magnified hologram was recorded by a detector placed approximately 5 m behind the sample. The hologram encodes the local phase shift, or equivalently the projected electron density distribution of the object. By choice of the focus-to-sample distance *x*_01_, which is typically a few millimeters to centimeters, the geometrical magnification 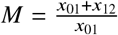 with sample-detector distance *x*_12_ and the effective pixel size 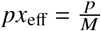 are adjusted, as well as the field of view.

### Data acquisition and image analysis

#### Scanning SAXS measurements

For the scanning SAXS experiments, the samples are continuously scanned by a piezo motor. Positioning of the sample in the focal spot is facilitated by an on-axis microscope. In view of radiation damage a ‘diffract before damage’ strategy is adopted (21):

1. position a pristine cell or pristine cellular region in the focal spot,
2. record a diffraction pattern with an exposure time, for which the signal is unaffected by structural damage,
3. move to a new cell/spot.

Regarding (2), the critical dose for damage is estimated from a test involving a variation of accumulation time, or from fractionating the dose over several sub-frames. This strategy is based on the tested assumption that the signal is stable up to a critical time depending on dose, dose-rate and resolution. This is the case for a pristine spot. Since previous exposures of neighboring scan points can induce a ‘bystander effect’, the step size is a further important parameter and has to be chosen sufficiently large (21). Therefore, this scheme results in rather coarse-grained real space maps of the cells. Tab. 2 presents the data acquisition parameters for the three different sample preparations. Data analysis was performed using the MATLAB nanodiffraction toolbox developed in our institute (46).

**Table 2:** Data acquisition parameters of the SAXS experiments.

|  | living | fixed | freeze-dried |
| --- | --- | --- | --- |
| fixative | - | KS | formaldehyde |
| step size ($\mu\text{m}$) | 5 | 10 | 2 |
| step size ($\mu\text{m}^2$) | $200 \times 200$ | $180 \times 180$ | $200 \times 200$ |
| stitching | 1 | $5 \times 5$ | 1 |
| acquisition time | 0.1 s | 0.1 s | 0.05 s |

#### X-ray holography

As a full field technique, x-ray holography does not require any lateral scanning of the sample, which makes it ideally suited for tomography. However, before any tomographic scan, the signal level, i.e. the contrast of the hologram has to be evaluated and the geometry and detector configuration has to be optimized. An overview of the different configurations of the beamline setup and the sample preparation tested in this work, is summarized in Tab. 3. Since contrast depends on the variations in electron density, freeze-dried cells result in much higher contrast than cells in solution. For CM recordings, we compared different detectors. We performed scans with a single-photon counting pixel detector (Eiger 4M, Dectris Ltd, Baden-Daettwil, Switzerland), offering highest dose efficiency due to photon counting. However, the large pixel size *px* = 75 *μ*m requires the sample to be moved very close to waveguide to reach high magnification, impeding tomographic rotations at high magnification. We therefore then turned to indirect detection based scintillator-based fiber-coupled sCMOS detectors. To this end, we used a single crystal scintillator (LuAG, thickness 20 *μ*m) with an sCMOS (Hamamatsu Photonics K.K., Hamamatsu, Japan) coupled by a 1:1 fiber optic plate, as well as another 1:1 fiber-coupled scintillator-based sCMOS camera (2048 × 2048 pixels, Photonic Science, Sussex, UK) with a custom 15 *μ*m thick Gadox scintillator with pixel size of *px* = 6.5 μm. This configuration was favored, since the whole cardiomyocyte fitted into the FOV and at the same time the effective pixel size was sufficiently small, namely *px*_eff_ = 45 nm at *M* = 143. The sample preparation and environment was also tested in view of compatibility with holographic tomography (holo-tomography). As described above, we tested hydrated unstained CMs as well as CMs with 1% OsO_4_ or PTA staining embedded in agarose. All cases resulted in a rather weak signal. For unstained cells embedded in agarose, almost no signal could be detected at the given photon energy. Sample mounting was another concern. As isolated CMs are not attaching to a substrate, one easily encounters drift during scan. For these reasons, we decided in this work to perform tomography scans only on freeze-dried CMs mounted on thin foils made from silicon nitride (Silson Ltd.). It is to mention, that for these experiments the size and shape of the SiN-frames is a compromise between the caused missing wedge, the tolerable tilt angle and the window’s stability: Thick and broad frameworks lead to a rather stable window at the cost of a large missing wedge. Windows with large edge lengths diminish the maximum tolerable tilt angle of the sample in horizontal and vertical direction due to the small *x*_01_ distance as required for high resolution cone beam recordings. Also the cell density must be chosen relatively low, otherwise neighboring cells will enter the field-of-view upon rotation and disturb the reconstruction of the target cell.

**Table 3:** Acquisition parameters of the x-ray holography experiments.

|  | I | II | III | IV |
| --- | --- | --- | --- | --- |
| preparation | freeze-dried | freeze-dried | freeze-dried | hydrated, stained |
| detector | Eiger 4M | sCMOS (20 $\mu\text{m}$ LuAG) | sCMOS (15 $\mu\text{m}$ Gadox) | sCMOS (15 $\mu\text{m}$ Gadox) |
| $px$ ($\mu\text{m}$) | 75 | 6.5 | 6.5 | 6.5 |
| $x_{01}$ (mm) | 20.8 | 20.8 | 35 | 35 |
| $x_{02}$ (mm) | 5413 | 5413 | 5011 | 5011 |
| $px_{\text{eff}}$ (nm) | 300 | 24 | 45 | 45 |
| photon energy (keV) | 8 | 8 | 7.5 | 7.5 |
| acquisition time (s) | $10 \times 0.1$ | 1 | 1 | 1 |

#### X-ray tomography

The configuration of the tomography setup as well as the sample preparation was chosen after the evaluation of the 2D holographic data sets. In order to avoid lack of contrast and motion artifacts, which would influence the quality of reconstructions, we focused on freeze-dried cells for tomographic recordings. At the same time, the size of single CMs (typically around 150 *μ*m) prompted us to choose a moderate magnification resulting in a pixel size of *px*_eff_ = 45 nm. To perform a proper phase reconstruction, projections were taken for four slightly different propagation distances (see Tab. 4) for each angular position. For each distance, 720 equidistant angular positions were recorded over 180°. At the beginning and the end of the tomographic recording, a set of 50 empty beam images were recorded in order to characterize the probing wave front. After the scan, 20 images of the camera dark current were acquired. The acquisition parameters of the experiment are summarized in Tab. 4.

**Table 4:** Acquisition parameters of the x-ray tomography experiments.

| parameter | value |
| --- | --- |
| photon energy $E$ (keV) | 7.5 |
| source-detector-distance $x_{02}$ (mm) | 5011.4 |
| source-sample-distance $x_{01}$ (mm) | {35, 36, 39, 45} |
| magnification $M$ | {143, 139, 128, 111} |
| pixel size $px$ ($\mu\text{m}$) | 6.5 |
| effective pixel size $dx_{\text{eff}}$ (nm) | {45.40, 46.69, 50.58, 58.37} |
| number of projections over $180^\circ$ | 720 |
| number of empty beam projections | 50 |
| number of darks | 20 |
| acquisition time (s) | 1 |

#### Phase retrieval and tomographic reconstruction

The phase and amplitude of the exit wave is reconstructed from the recorded holograms by phase retrieval, either based on iterative projection algorithms or in some cases direct Fourier-filtering, which enables to obtain a phase or projected electron density map of the cell. Note, that for a biological cell at the given photon energy, absorption can be neglected, and the cell can essentially be regarded as a pure phase object. In this study a nonlinear approach of the contrast transfer function (CTF) was used for phase retrieval (47, 48). The phase distribution in the object plane *ϕ*(**r**_⊥_) retrieved by the linear form of the CTF is given by

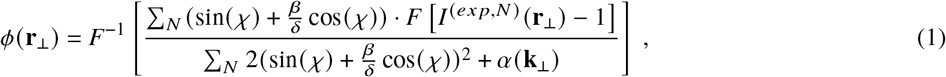

with natural units 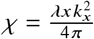 for the (squared) spatial frequencies, measured intensities in the detector plane *I* ^(exp)^(**r**_⊥_), 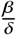 ratio of imaginary and real part of the refractive index *n*, N indicated the index of the respective distance and a frequency-dependent regularization parameter *α*(**k**_⊥_) (lim_1_ for high and lim_2_ for low spatial frequencies). The nonlinear approach uses the solution of the CTF as input for an iterative phase retrieval based on a minimization of the Tikhonov functional (47). Thus, the accuracy of the reconstructed phase is not less than the results of the CTF. The parameters of the phase retrieval are given in Tab. 5.

**Table 5:**
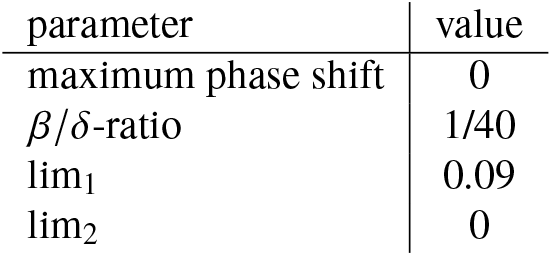
Phase retrieval parameters for the nonlinear Tikhonov approach of the CTF for the x-ray tomography experiments.

Before tomographic reconstruction, an alignment of the single projections based on the linogram was performed to correct for a vertical movement of the sample. A ring removal was not necessary since artifacts from oversensitive pixels were reduced from the previous shifting based in the alignment of the linogram. For tomographic reconstruction the ASTRA toolbox was used (49, 50). Projections in the range where the frame of the SiN window enters the FOV were excluded.

After tomographic reconstruction, the local electron density can be computed from the phase shift induced by each voxel length *v* as

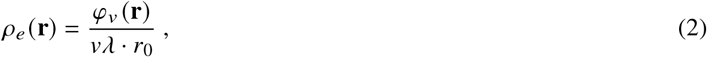

with wavelength *λ* and *r*_0_ the classical electron radius.

#### Dose calculation

The dose was calculated using (51, 52)

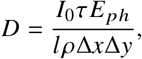

with the total photon flux *I*_0_, the exposure time *τ*, the energy *E*_*ph*_ and the size of the illuminated area Δ*x*Δ*y* as listed in Tab. 1. Cellular samples are commonly described by H_50_C_30_N_9_O_10_S. For the used experimental parameters this yields a mass density 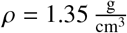 and a mass attenuation length *l* = 7.46 · 10^−4^ m (51, 52). The choice of image acquisition parameters for diffraction experiments was guided by our previous investigation and simulation of radiation damage (21). The average dose per shot in the diffraction experiments was *D* = 5.3 · 10^5^ Gy. For the holo-tomography, the total dose for a tomography scan with 720 projections an acquisition time of 1 second was estimated to *D* = 1.1 · 10^5^ Gy, where the illuminated area Δ*x*Δ*y* is given by the effective area (FOV) covered by the detector. This shows that the dose of a complete tomographic scan is still lower than a single shot of the diffraction measurements.

## RESULTS AND DISCUSSION

### 2D holography of isolated CMs

Figure 2 shows single projections for different sample preparations, detectors and magnifications, which were acquired to test contrast and stability imaging conditions. Table 3 summarizes the parameters tested for the different sample preparations and beamline configurations. Additionally, an intensity profile highlights the interference fringes originating from phase effects caused by the interface of the cell and the surrounding medium is marked by the red lines in the image. Noise was reduced by averaging the intensities from pixels perpendicular to the lines corresponding to a linewidth of 1 *μ*m. The fringes originating from changes in electron density indicate the holographic nature of image formation, and contain structural information. In Fig. 2a a projection of a freeze-dried CM recorded with a single photon counting detector (Eiger 4M) at an effective pixel size of *px*_eff_ = 300 nm is shown. In this projection the interface between cell and surrounding air dominates the contrast. Besides that, the position of the nucleus can be determined and the signal of the sarcomeric structure starts to appear. For this configuration, the field of view (FOV) is not limited by the detector size (FOV = *px*_eff_ · *N*_pixel_) but by the illuminated area of the waveguide. Only 351 × 351 pixels of the detector are shown in the image. In order to go to higher magnifications, the sample-detector distance has to be increased. Since the detector distance cannot exceed the length of the experimental hutch, the sample has to be moved closer to the wave guide. This would be possible for 2d holographic measurements, but since it is the goal to image the 3d structure and therefore the sample has to be rotated, the minimal source-to-sample *x*_01_ distance is limited by the size of the SiN frame on which the CMs are mounted.

**Figure 2:**
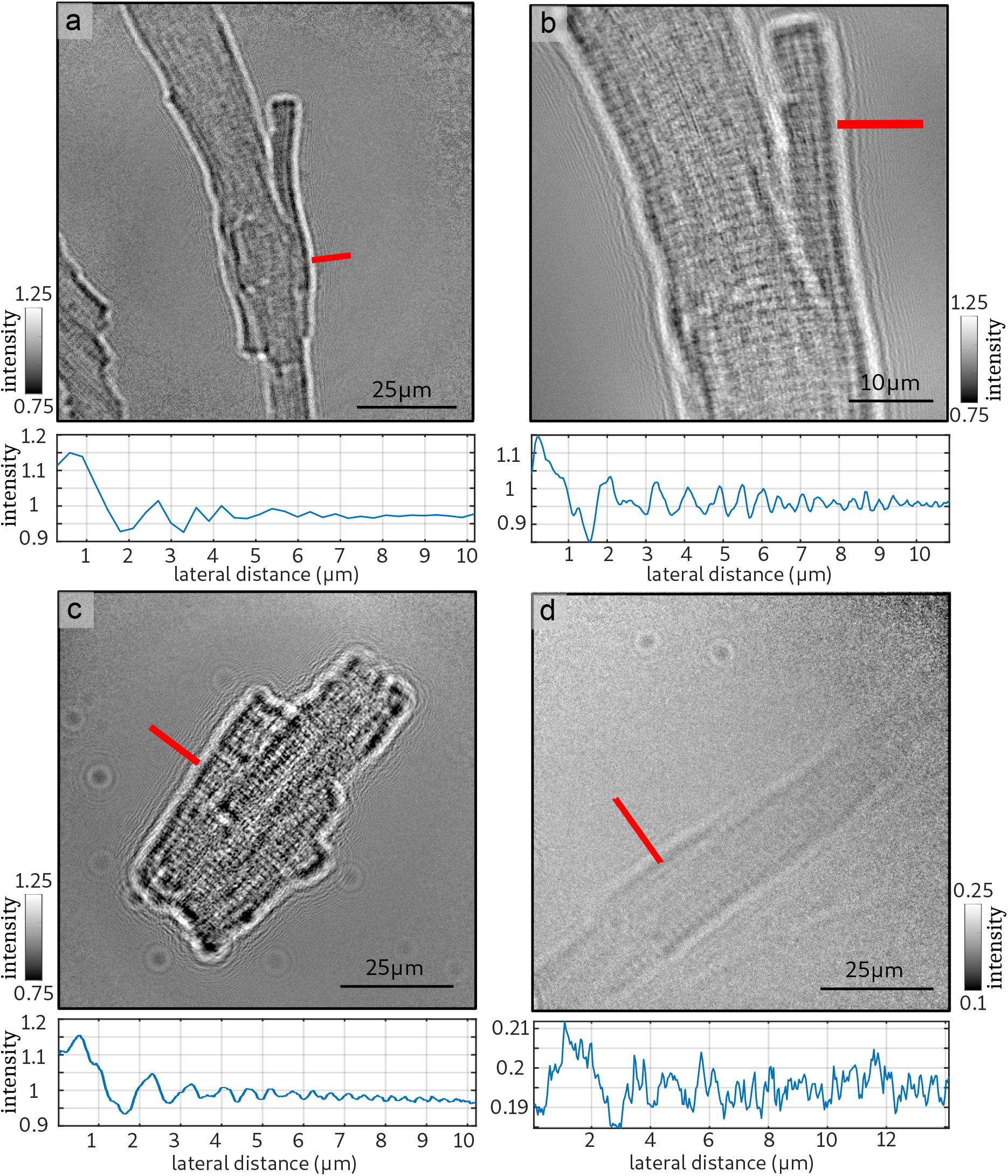
Exemplary projections of the 2D holography measurements. To optimize the imaging conditions, different detectors, magnifications and sample preparations were tested. The red lines mark the area of the intensity profiles shown below each projection. (a) Projection of a freeze-dried CM using a single photon counting detector (Eiger 4M) resulting in an effective pixel size of *px*_eff_ = 300 nm. (b) Projection of the same CM using a fiber coupled sCMOS Camera (20 *μ*m LuAG) with *px*_eff_ = 24 nm. (c) Projection of a freeze-dried CM at an effective pixel size of *px*_eff_ = 45 nm (15 *μ*m Gadox) (d) Projection of a CM stained with 1% OsO_4_, embedded in agarose and mounted in a polyimide tube with a diameter of 1 mm.

Figure 2b shows a projection of the same cell acquired with a fiber coupled sCMOS Camera (LuAG) at an effective pixel size of *px*_eff_ = 24 nm. In this case the entire detector is illuminated and smaller effective pixel sizes can be achieved. The signal from the cellular membrane and also sarcomeric structure is much stronger than in the projection at lower effective pixel size. The fringes around the CM are very clear as also indicated by the profile shown below the projection. At this high magnification only a part of the cell can be imaged due to the relatively small FOV. Further, thermal drift of the sample and vibrations of the setup in the range of a few manometers led to artifacts in the tomographic reconstruction. Fig. 2c shows a projection of a freeze-dried CM was acquired at moderate effective pixel size of 45 nm using the sCMOS camera with a 15 *μ*m Gadox scintillator. In this case an entire CM could be imaged within a single projection and the signal emerging from the cell-air interface as well as from the sarcomeric structure can be identified. Since the goal is to image an entire CM we decided to use this beamline configuration for the x-ray tomography.

In a next step, the CM in hydrated condition were investigated. Figure 2d shows a projection of a CM stained with 1% OsO_4_, embedded in agarose and mounted in a polyimide tube with a diameter of 1 mm. The image was acquired with the same beamline configuration as for the freeze-dried cell shown in Fig. 2c. In this case, the signal from the cell is very weak. Note that the quality of the signal is also diminished by the strong absorption from the agarose in the 1 mm tube. In view of the above, tomographic scans were recorded for the freeze-dried sample with the sCMOS (15 *μ*m Gadox) camera at an effective pixel size of 45 nm.

### 3D structure of a single cardiomyocyte

The 3D structure of the cardiomyocyte was reconstructed by propagation-based phase-contrast tomography in the holographic regime. Figure 3 shows the results of the 3D cellular structure. For all projections phase retrieval was performed by the nonlinear Tikhonov approach of the CTF as explained above. An example for a projection and the corresponding reconstruction by phase retrieval is shown in Fig. 3a. The sarcomeric structure is clearly visible. The brighter stripes with a lower phase shift, perpendicular to the orientation of the cell correspond to cellular structures with a lower electron density and probably indicate the M-line of the sarcomere. Based on the phase retrieved projections, tomographic reconstruction was performed. Figure 3b shows a volume rendering of the entire CM. The total volume of the cell was determined to about 16 200 *μ*m^3^. A slice of the reconstructed 3d electron density is shown in Fig. 3c. Sub-cellular structures of the CM are resolved. In the case of this cell, we have two nuclei instead of just one nucleus. Both are located in the center of the cell. Further, the myofibrils and the sarcomeric structure can be identified. Between the elongated myofibrils small dense organelles with a diameter of roughly 500 nm are visible. As justified further below, these structures can be attributed to mitochondria, which are known to be abundant in CMs, providing the required energy for force generation/contraction. They are arranged in chains next to the myofibrils. Within the range of one sarcomere the mitochondria frequently appear in pairs of two elongated blobs. Figure 3d shows a maximum projection of the 3d confocal microscopy stack. The actin skeleton was fluorescently labeled and the orientation and position of the elongated actin structures indicate that fibers of 3d electron density correspond to the actomyosin filaments (myofibrils) of the CM.

**Figure 3:**
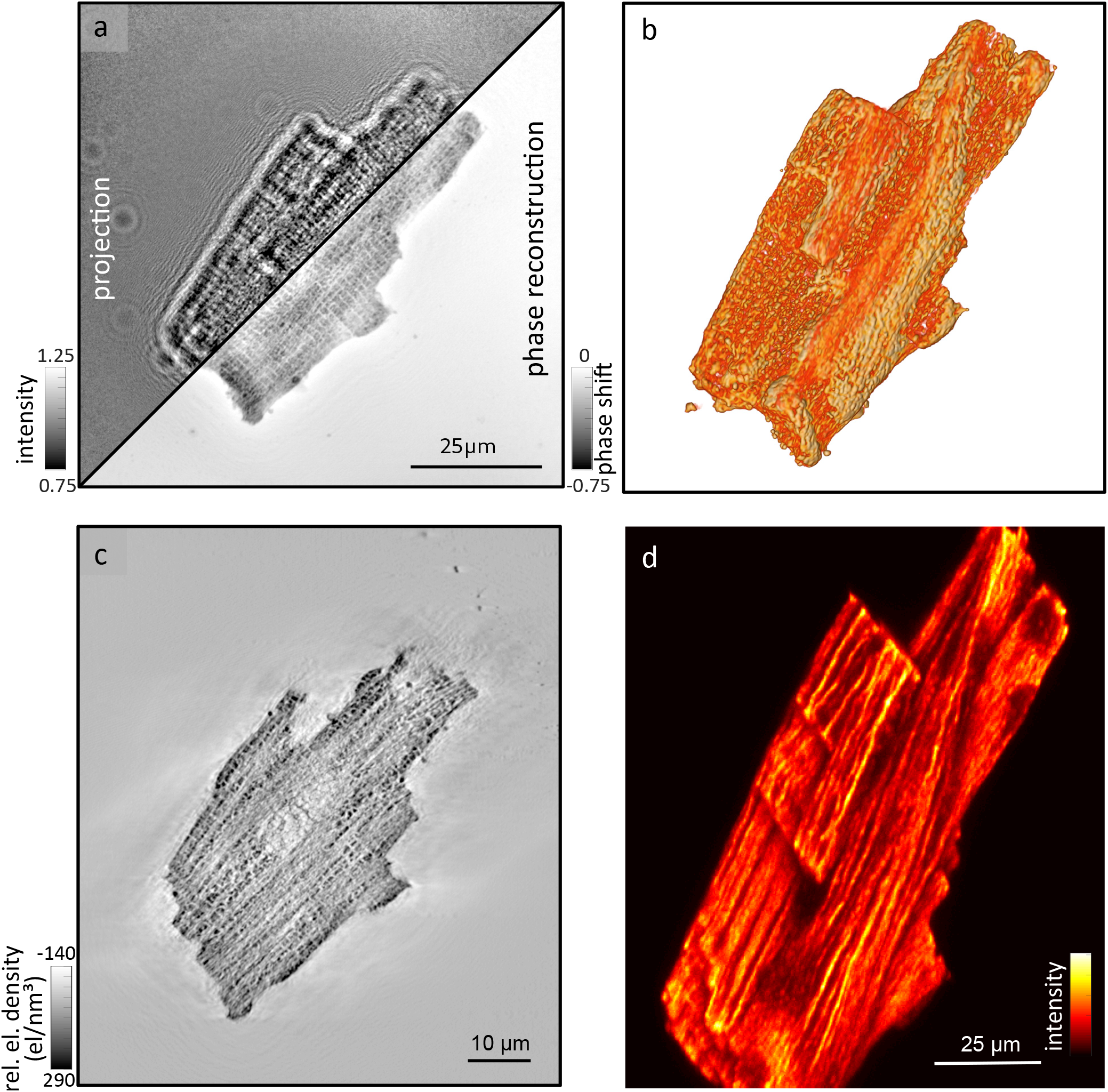
Phase retrieval and reconstruction. For each of the 720 angular positions, phase retrieval was performed based on the projections acquired at different Fresnel numbers. An exemplary projection and the corresponding phase retrieval is shown in (a). The 3d structure was reconstructed by the inverse radon transform. A volume rendering of the CM is shown in (b). A 2d slice though the reconstructed volume is shown in (c). Sub-cellular structures such as the nuclei, mitochondria and myofibrils can be identified. Further, the fluorescently labeled actin skeleton of the same cell was imaged by a confocal scan. A maximum projection of the microscopy stack is shown in (d).

### Segmentation of mitochondria

The dense structures observed in the electron density can be attributed to mitochondria for several reasons. First, mitochondria exhibit a high electron density close to that of a protein crystal (53) and therefore appear as dark structures compared to the surrounding actomyosin in the reconstruction. Second, they are of elliptical shape with a diameter in the range of 0.5-1 *μ*m (54). Third, they are located next to the elongated myofibrils, as already known from histological observations and 2d electron microscopy images of healthy cardiomyocytes (55). Most importantly, apart from the actomyosin which appears in form of the expected sarcomeric and myofibrilar structure in the reconstruction, there is no equally abundant organelle in CMs which could explain this additional structure.

The segmentation of the mitochondria was performed using a software designed for the analysis of 3d (fluorescence) microscopy and tomography data sets (Vision 4D, arivis AG, Munich, Germany). To this end, the 3d volume from the x-ray tomography analysis was converted to a 8 bit .tif stack and loaded into the program. Using a ‘blobfinder’ tool, roundish structures can be identified. For the present data set a characteristic size of 12 px ≙ 520 nm was chosen, which fits well to the size of murine mitochondria. After this step more than 45 000 blobs, which could be associated to mitochondria, parts of the sarcomeres, membrane residues and other sub-cellular compartments, were tracked. Next, false positive blobs were removed by thresholding the minimal volume to a minimal voxel count of 500 voxels per detected volume. This volume corresponds to a perfect sphere with a radius of 5 px (225 nm). Further selection of the mitochondria was done by a threshold in density. Only blobs containing voxels with at least 100 electrons/nm^3^ were included. As a result of this analysis pipeline, 14 334 blobs matching the size and position of the mitochondria remained, which corresponds to an estimated average density of about 0.88 mitochondria per *μ*m^3^.

Figure 4 shows the results of the segmentation. The mitochondria appear as small dark spots in the slice of the reconstructed electron density shown in Fig. 4a. The red box marks an area for which the quality of the segmentation is shown in Fig. 4b. The segmented mitochondria of this slice are shown in random color, in order to facilitate to distinguish between neighbors. In the area at the bottom of the slice, two mitochondria can be assigned for each sarcomere (from M-line to M-line). Based on this segmentation of the entire 3d volume, the electron density of all mitochndria was analyzed. A histogram of the extracted electron density is shown in Fig. 4c. From this distribution a mean electron density of freeze-dried mitochondria was determined to 162 el/nm^3^. Further, the size of the mitochondria can be quantified by the length of the segmented volumes along its main axis. The short, middle and long side of segmented blobs are shown in Fig. 4d. From this distribution, the diameter and length of the elipsoid-shaped mitochondria can be extracted. The peak in the histogram of the short axis (blue) corresponds to an diameter of 585 nm, while the long side (red) has a peak for a length of 1125 nm. For the short side a small increase can also be seen at about 1200 nm, and also for the medium and long axis an increase in the range of the double length of the peak can be identified. This increase can be explained by a pooling of neighboring mitochondria as one selection. Thus, we conclude that the total number of mitochondria is underrated. The mean volume of a mitochondrium was determined to 0.23 *μ*m^3^. The smaller size of the mitochondria may result from a volume change at the mitochondrial level during the sample preparation. The volume fraction was determined to about 20% by comparing the sum of mitochondria volume to the total cell volume. Considering the shrinking of the mitochondrial level by a factor of approximately 2, these results are consistent with a volume fraction between 29% and 36% in the literature. Besides analysis of density, size and shape of mitochondria, this analysis yields the location of the mitochondria in the cell.

**Figure 4:**
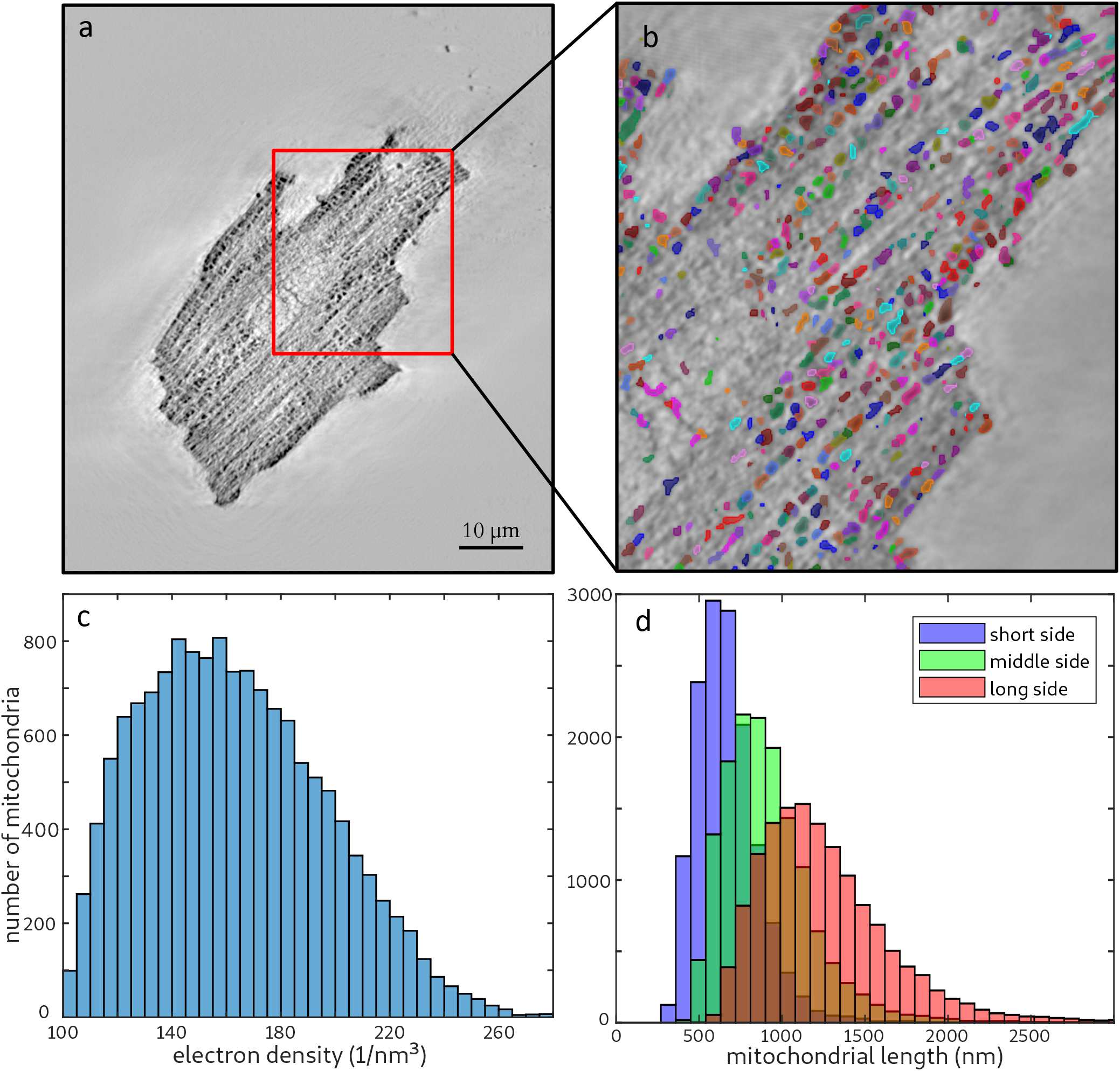
Mitochondria were segmented based on the reconstructed 3d electron density of the CM. We used an image processing software (Vision 4D, arivis AG, Munich, Germany) developed for microscopy data. The dense roundish mitochondria were identified by a ‘blob-finder’ algorithm and a threshold in size and density. In (a) one slice from the tomographic reconstruction is shown. In order to visualize the quality of the data analysis a magnification of the area marked with a red box and the corresponding segmentation is shown in (b). The segmentation is colorized in a random color to distinguish neighboring mitochondria. Based on the segmented data from the entire volume further analysis can be performed. In (c) a histogram of the maximal electron density of the segmented mitochondria is shown. (d) Distribution of the short (blue), medium (green) and long (red) axis of the segmentation. From this information the diameter and length of an average mitochondrium can be calculated.

### Characterization of the actomyosin lattice

The molecular structure of heart tissue cells was analyzed by scanning x-ray diffraction. Figure 5 shows a typical result for scanning diffraction from freeze-dried CMs, using a microfocus beam. We include this example for comparison with the live recordings below. As becomes immediately evident, the x-ray darkfield signal is particularly strong in this case. Furthermore, freeze-dried CMs can be imaged with relatively high sampling in real space, since less free radicals are produced in the absence of water/buffer. More importantly, free radicals do not spread by diffusion as in hydrated environments (21). In Fig. 5a, an optical micrograph of (fluorescently labeled) freeze-dried CMs taken by the beamline on-axis video microscope (OAV) is shown. The cells are clearly visible in bright field contrast. Using the live images of the OAV, the samples were aligned for the x-ray diffraction measurements. An example for an x-ray darkfield image from the analysis of freeze-dried cells is shown in Fig. 5b. Pixels with an intensity below 1.5 · 10^6^ counts were masked in white. The step size was 2 *μ*m, the acquisition time 0.05 s. In Fig. 5c, an isolated diffraction pattern is shown, representing a single shot from the freeze-dried cells. There are no actomyosin reflections visible in the diffraction pattern, indicating that the structure of the cells is damaged by freeze drying. The diffraction pattern shows an anisotropy which makes it nevertheless possible to determine the local actomyosin fibril orientation. The fiber orientation and anisotropy, shown in Fig. 5d, is obtained from an automatized principal component analysis (PCA) in the same way as in (20, 46). In order to improve the quality of the diffraction data, we turn to recordings of hydrated and living cells using a polypropylene chamber. In view of radiation damage, a ‘diffract before damage’ strategy is adopted (21).

**Figure 5:**
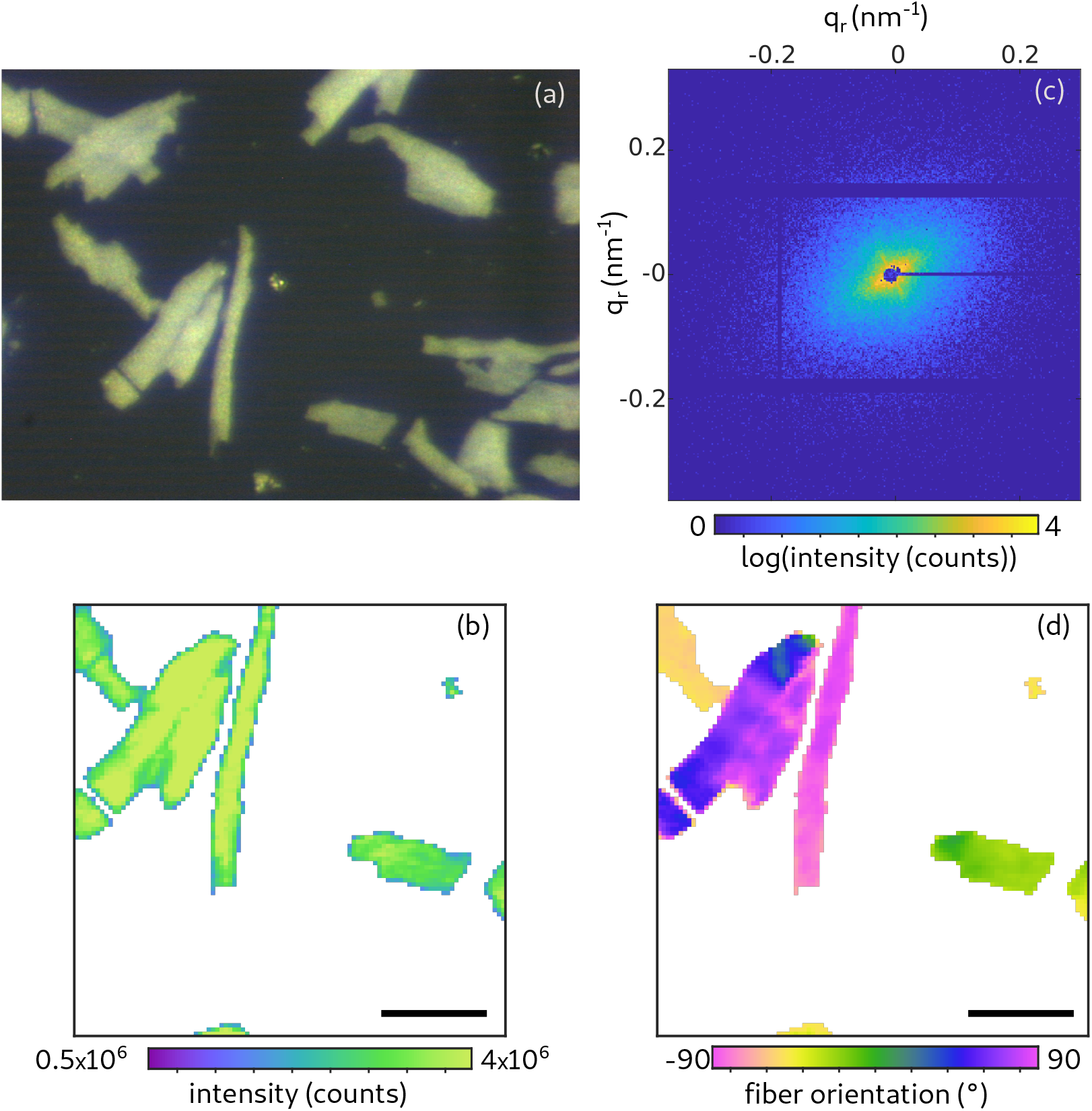
Scanning diffraction from freeze-dried CMs. (a) Micrograph recorded with the beamline on-axis optical microscope (brightfield). (b) X-ray darkfield image. The image is formed by integration of all detector counts (outside the beamstop), i.e. the integrated scattering signal. Pixels with an intensity below 1.5 · 10^6^ counts are masked in white. Scale bar: 50 *μ*m. (c) Isolated diffraction pattern of a single shot on a freeze-dried cell (0.05 s accumulation time). (d) Orientation of the acto-myosin fibrils.

Figure 6 illustrates the data collection and processing, which closely followed (22), but now applied to living cells. We also took into account the constraints of sparse sampling and reduced dose to collect a signal which is not spoiled by radiation damage (21). For data analysis, the nanodiffraction-toolbox (46) was used. In Fig. 6a the liquid chamber used for the measurement of living CMs is shown. Figure 6b shows the example of an isolated diffraction pattern recorded by a single shot. The acquisition time was 0.1 s. In contrast to the freeze-dried CMs, the signal of the living CM shows clearly both the (1,0) and the (1,1) reflection in the diffraction pattern. The cells were raster scanned with a step size of 5 *μ*m, yielding a coarse darkfield map of the scanned CMs. An example for such a map can be seen in Fig. 6d. Note, that the background in the map is colored in white based on a threshold of minimum photon counts. Next, each diffraction pattern above this threshold was azimuthally averaged and a mean background was subtracted to obtain a local SAXS curve. The model function

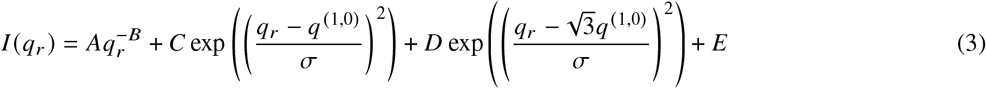

was fitted to this data, where *A − E* and *q*^(1,0)^ are fit parameters. While 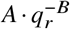 describes a monotonous SAXS contribution, the actomyosin reflections are modeled as Gaussians with amplitudes C and D and positions *q*^(1,0)^ and *q*^(1,1)^. Note that the relation 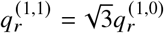 was used to obtain the position of the (1,1) reflection. The reflection width was set to *σ* = 0.0184 nm^−1^. An example of this fit for the isolated shot can be seen in Fig. 6c. This fit yields the position of the (1,0) position. In Fig. 6f the *q* ^(1,0)^ reflection positions of the sample are plotted. Using PCA, the fiber orientation was determined for the diffraction pattern for all scan points. An example for the fiber orientation is shown in Fig. 6e.

**Figure 6:**
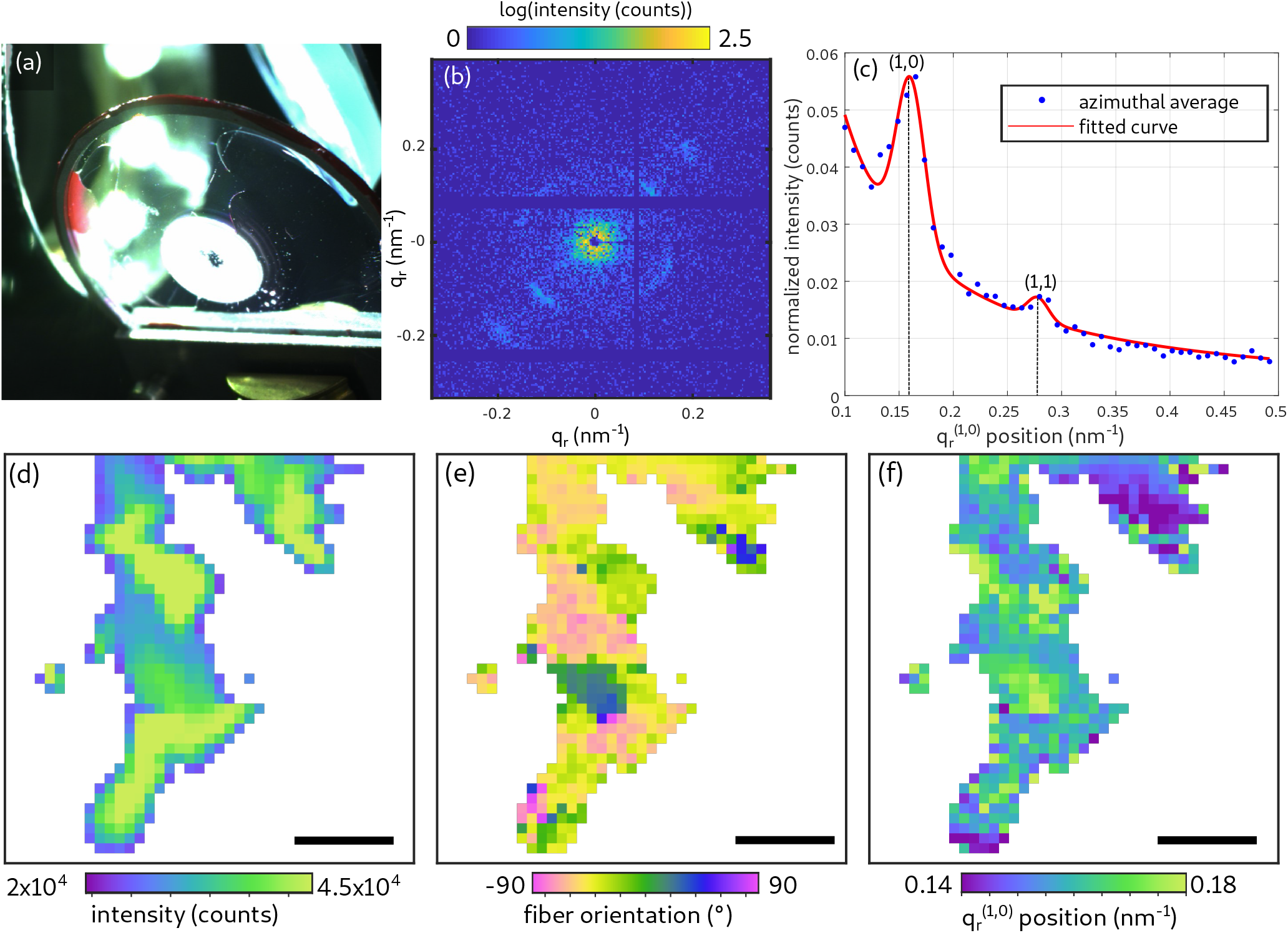
Signal processing of scanning diffraction data: living CMs. (a) Photograph of the sample chamber used for measuring living CMs. (b) Diffraction pattern of a living muscle cell. The (1,0) and the (1,1) reflection is visible in the pattern. (c) 1d azimuthal average of the diffraction pattern on the left. The fitted curve was obtained using the fitting function introduced in Equation 3. (d) Darkfield map of a cluster of living cells. Scale bar: 50 *μ*m. (e) Orientation of the myosin fibers obtained by performing a principal component analysis. (f) Position of the (1,0) reflection obtained from the fitted curves. In (c)-(e) all pixels, with a darkfield intensity below 2 · 10^4^ counts were masked in white.

For comparison to the living CM, we also investigated chemically fixed, hydrated cardiac tissue cells. Figure 7a shows an example for a darkfield map of such a recording. Multiple scan areas covered by continuous piezo scanning, each with a size of 200 × 200 *μ*m^2^ and an acquisition time of 0.05 s per scan point, were stitched using a stepper motor below the piezo. The entire composite with 5 × 5 piezo scans results in the darkfield map in Fig. 7a. The different scans areas are indicated by boxes. The acquisition time for each scan point was 0.05 s. In Fig. 7b, a zoom of the marked box in (a) is shown. In Fig. 7c the calculated fiber orientation of this region is plotted. The data shows, that the fiber orientation throughout a cell is roughly constant. An example for an azimuthal average of a single diffraction pattern of a chemically fixed cell is shown in Fig. 7d. Only the first peak representing the (1,0) reflection is visible. This was the case for all shots recorded from all chemically fixed cells. Therefore, the signal of the (1,1) reflection was excluded for the fit of the fixed cells. The results for the q_r_^(1,0)^ position for one region of the map shown in Fig. 7e.

**Figure 7:**
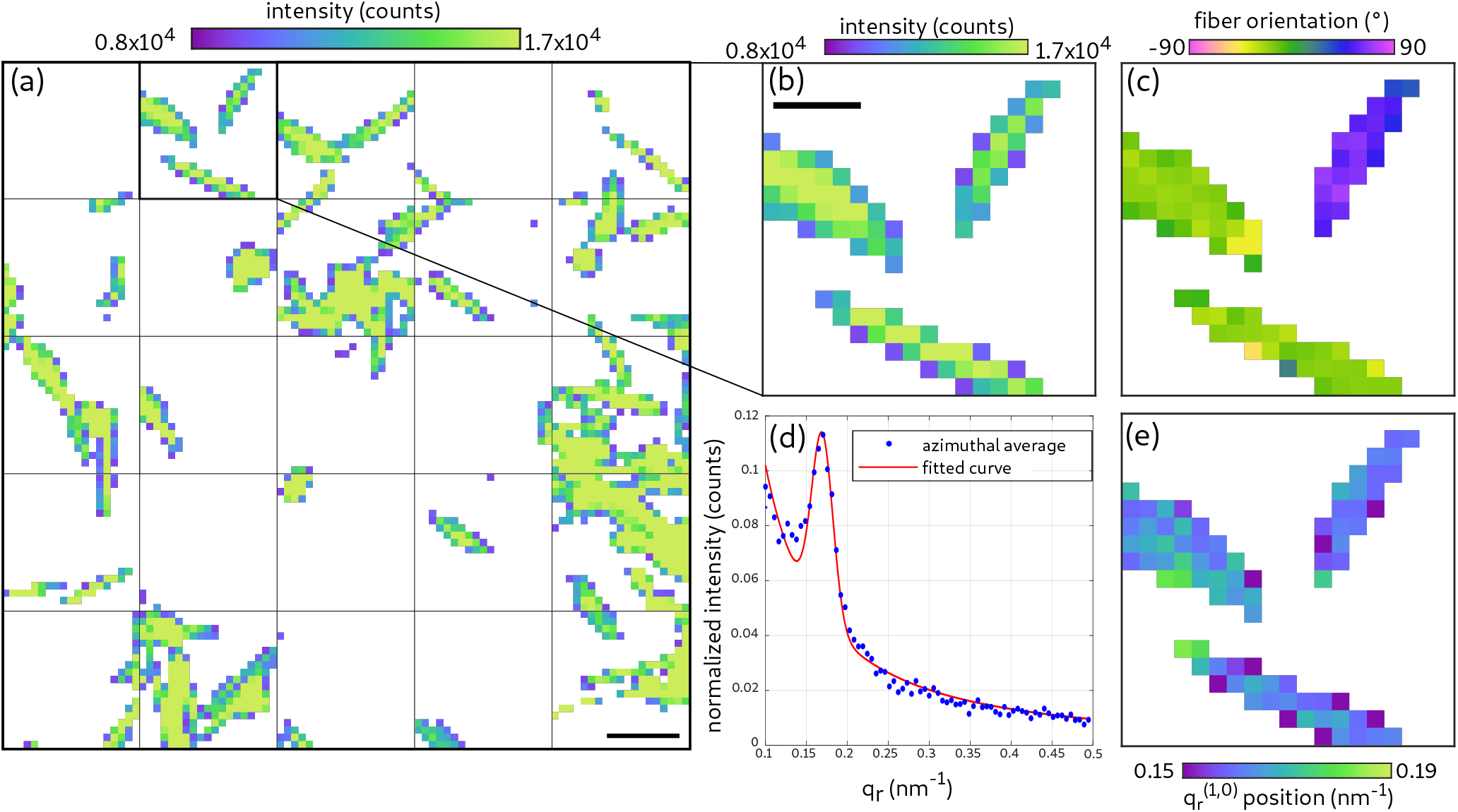
Scanning diffraction: hydrated and chemically fixed CMs. (a,b) Darkfield maps of isolated, fixed cardiomyocytes. Scale bar: (a) 100 *μ*m, (b) 50 *μ*m. (c) Orientation map of the actomyosin fibrils. (d) 1d azimuthal average of a single shot from a fixed cardiomyocyte. In contrast to Fig. 6 (c) only the (1,0) reflection is visible. (e) Position of the (1,0) reflection. All pixels with a darkfield intensity below 0.8 · 10^4^ counts were masked in white.

Next, we turn to the recordings of living cells, which are displayed in Fig. 8. These cells were in an initially living state as one could derive from their contraction inside the sample chamber before the measurement. However, one has to keep in mind the challenging issue of radiation damage arising during data acquisition which will have a considerable effect on cellular structures. Thus, we can only claim that the cells are living initially at the beginning of the scan or when recording shots from a new cell. In Fig. 8a a darkfield map of a cluster of living CMs is depicted; the same which was already shown before when explaining the workflow. The boxes marked in in the map of the darkfield signal are represented in detailed view in Fig. 8b-d. For the three areas, the amplitude of the first peak representing the (1,0) reflection is shown in (1). The first peak was visible in all observed boxes. On the contrary, this is not the case for the amplitude of the second peak representing the (1,1) reflection which is shown in (2). The harmful effect of radiation damage can be observed by the loss of (1,1) reflection signal which is visible in the first few lines of the first area (Fig. 8b.2). In this region, the signal of the (1,1) reflection is clearly visible in the first scan points, but then vanished. This observation is consistent with previous observations on muscle tissue, where the intensity of the second peak vanished when tissue was affected by radiation damage (21). Here we also observe that the intensity of the second peak decreases, indicating that living cells are strongly affected by radiation damage. This signal decrease is plausible, since the scan points were acquired row by row from top left to bottom right, and free radicals spread easily during the scan. Hence, for the first region, the concentration of free radicals was lower, and less time was available for their diffusion or for an apototic reaction of the cells. The map of the (1,1) peak amplitude also shows that the intensity decreases line-by-line.

**Figure 8:**
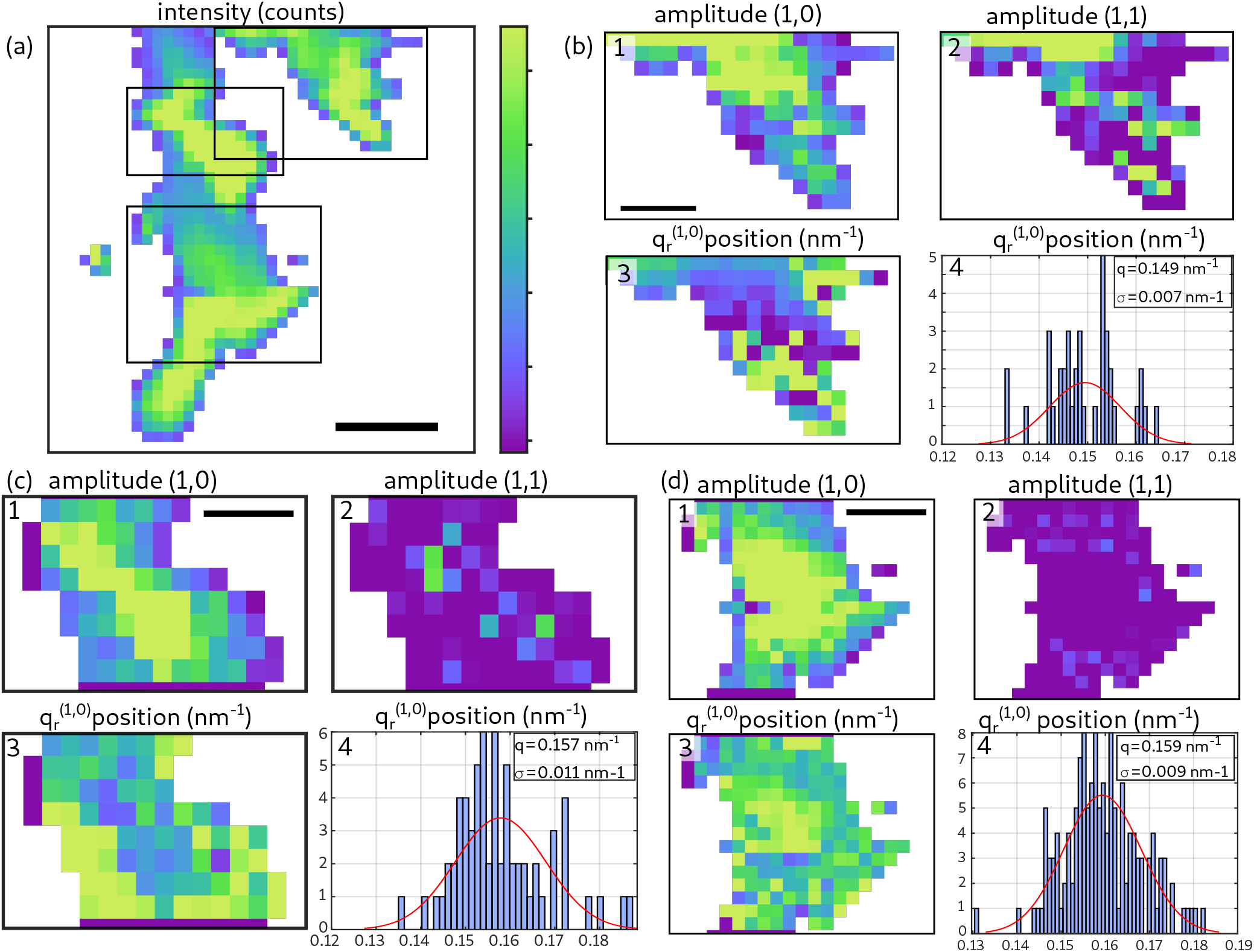
Scanning diffraction: living CMs. (a) Darkfield image showing three CM clusters magnified in (b,d and d). Scale bar: 50 *μ*m, colorbar: linear range 2 · 10^4^ - 4.5 · 10^4^ counts. In all figures pixels with a darkfield intensity below 2 · 10^4^ counts where masked in white. The structural parameters of the actomyosin lattice were calculated, namely 1. amplitude of the (1,0) reflection, 2. amplitude of the (1,1) reflection, 3. q_r_^(1,0)^ -lattice spacing and 4. histogram of the q_r_^(1,0)^ position. (1) Scale bar: 25 *μ*m, colorbar: linear range 0 - 0.1 counts/pixel. (2) Colorbar: linear range 0.5 · 10^−3^ - 4 · 10^−3^ counts/pixel. (3) Colorbar: linear range 0.14 nm- 0.18 nm^−1^. (4) In area (b) only shots where two peaks are visible, contributed to the histogram.

The q-spacing of the the actomyosin lattice was evaluated for the entire dataset. The subfigures with index (3) show the map for the q-spacing of the different areas. Boxes with index (4) show the distribution of the q-spacing (histogram). In Fig. 8b.4 only shots with two visible peaks were selected for the histogram. In Fig. 8c.4, and Fig. 8d.4 only shots with one peak contributed to the histogram. A Gaussian was fitted to every histogram to obtain the mean and standard deviation. The analysis of the histograms shows that the q-spacing for the first box (*q* = 0.149 ± 0.007 nm^−1^) is smaller than the q-spacing in box (c) (*q* = 0.157 ± 0.011 nm^−1^) and box (d) (*q* = 0.159 ± 0.009 nm^−1^). This leads us to the conclude that the q-spacing increases when the cells are affected by radiation damage. Since the presented scans were also partly affected by damage as inferred from the decrease of the (1,1) peak, we adapted our measurement strategy, and in addition to the scans also took diffraction pattern from isolated points within the CMs, moving to new cells for each shot.

Finally, the violin plot in Fig. 9 compares the distribution of the q-spacing for (initially) living, hydrated CMs and CMs, which were chemically fixed. In fixed CMs, only the (1,0) reflection was visible in the diffraction pattern. For the characterization of living cells, the diffraction patterns from the data already seen in 8, as well as additional single shots from freshly prepared samples were used. Only patterns with two visible peaks were used for the calculation of the lattice spacing of living CMs. The q-spacing determined for fixed cells is larger than the one found for the living cells. To quantify the mean and standard deviation, Gaussians were fitted to the distributions. The horizontal lines in the violin plot indicate the mean value. For living CMs a q-spacing of *q* = 0.148 ± 0.006 nm^−1^ and for chemically fixed CMs *q* = 0.167 ± 0.006 nm^−1^ was obtained. The corresponding actomyosin lattice spacing 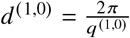 was 37.6 nm and 42.5 nm, for the fixed and living cells, respectively.

**Figure 9:**
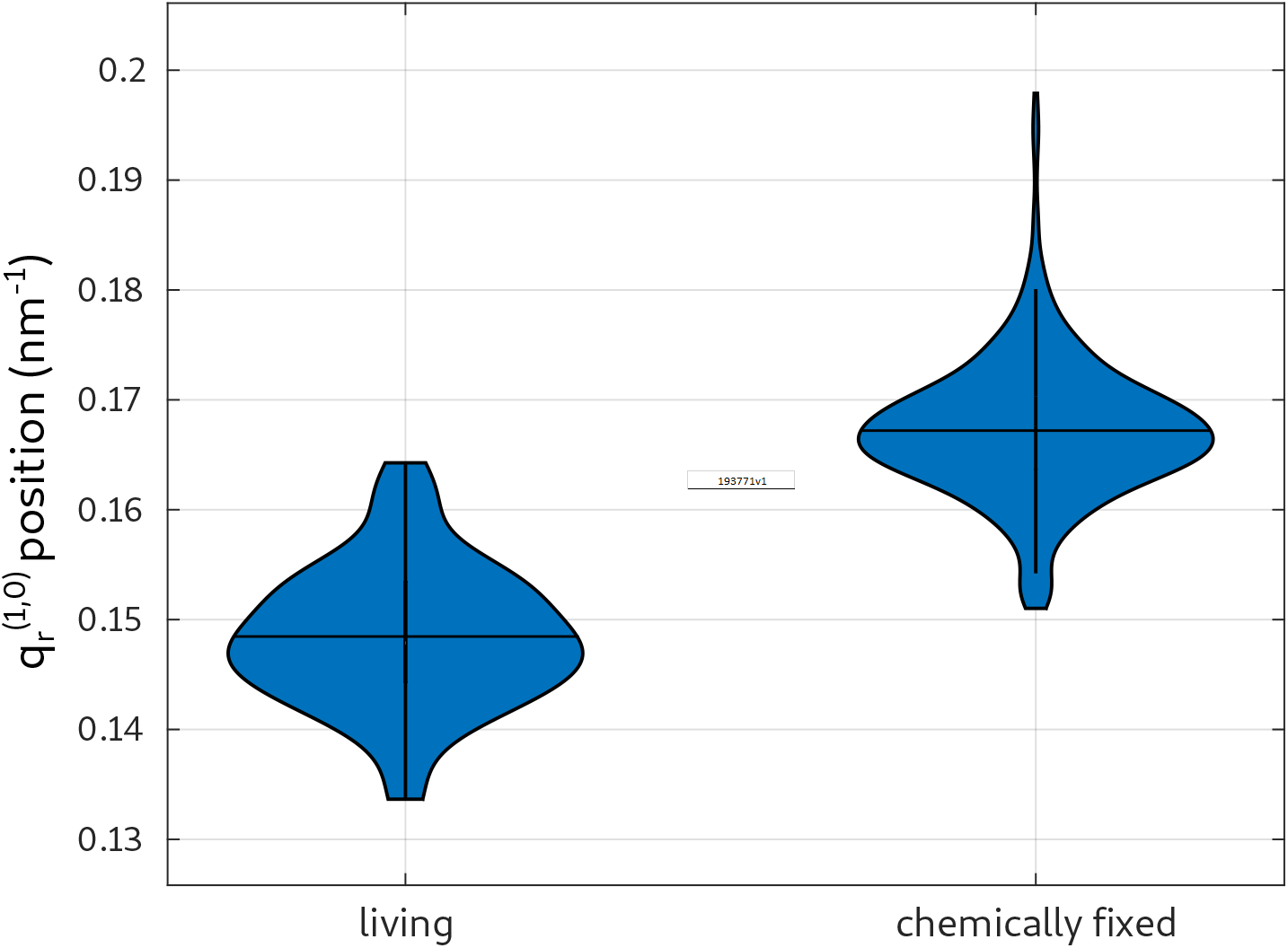
Actomyosin lattice spacings. Distribution of the lattice spacing obtained by the position of the q_r_^(1,0)^ reflection for living and chemically fixed CMs. The horizontal lines in the violin plot indicate the mean value of the q-spacing. The corresponding actomyosin lattice spacing determined to 37.6 nm for chemically fixed CMs and 42.5 nm for living cells.

## SUMMARY AND CONCLUSION

In this work, we have combined full-field 3d coherent x-ray imaging and scanning x-ray diffraction to analyze the cellular and molecular structure of isolated adult CMs of wild type mice. The 3d electron density distribution of a single CM was reconstructed from propagation based x-ray phase contrast tomography with a voxel size smaller than 50 nm. Single myofibrils, the sarcomeric organization and location of mitochondria were visualized within a single cell without sectioning. The molecular structure of the actomyosin lattice was probed by scanning x-ray diffraction, using CRLs for micro-focusing.

For the first time, the diffraction signal of the acto-myosing contractile unit could be observed for living cells. Importantly, while previous recordings of living cells only showed a monotonous decay of the SAXS signal (56), the diffraction signal of CMs shows characteristic peaks which can be fitted to determine the actomyosin lattice spacing. A comparison between living cell and chemically fixed cell recordings indicates that the characteristic lattice distances shrink by approximately 10% upon fixation, namely from 42.5 nm to 37.6 nm in the 1, 0 lattice planes.

Despite its merit in extending diffraction analysis to living CMs, the present study also indicated limitations of this approach, imposed by radiation damage and low signal-to-noise. For future work, it would be desirable to probe different states of contraction, eventually in combination with well controlled forces acting on the cell. To this end, only isolated shots in fresh spots should be collected, guided by online light microscopy, and correlated with the contraction state. Continuous replenishment of samples and washing out of free radicals could be achieved by suitable microfluidic devices. Finally, high throughput lattice parameter determination at the single cell level, eventually in form of ‘diffraction flow cytometry’ could be used as a diagnostic tool for dissociated tissue. To this end, genes regulating actomyosin structure in patient-derived stem cells would be targets of particular interest (57). Concerning the tomography, investigations of hydrated or even living cells seem very challenging based on our present results. Frozen hydrated (vitrified) cells after plunge freezing (58–62), however, could be a suitable approach to probe the native structure in 3d. In this study, we could for example determine an average density of about 0.88 mitochondria per *μ*m^3^ as well as the average size of a mitochondrium in such a CM. A number of technical improvements can also be foreseen: a small glass tube with thin walls and a hydrogel could provide stability for single cells and minimize absorption. The sample could be moved closer to source spot for higher magnification and single photon counting pixel detectors with higher quantum efficiency could be used.

Concerning the sequence of correlative imaging of the same cell, it is important to first perform optical fluorescence microscopy, second holo-tomography, and finally diffraction, in view of mitigating the radiation damage. The final state could then be probed again by optical microscopy to evaluate damage.

## AUTHOR CONTRIBUTIONS

TS and MR designed research. KT developed adult cardiomyocyte protocols and contributed expertise. MR and TS carried out the x-ray holography experiments. MR, CN, TS, JDN, carried out the diffraction experiments. JDN wrote analysis software. MR carried out the phase retrieval and tomographic reconstruction, CN, JDN carried out the SAXS analysis, supervised by TS.

MB and MR prepared the freeze-dried cells and recorded the florescent microscopy data. MR, TS, and CN wrote the article; all authors read final data.

## ACKNOWLEDGMENTS

We thank Dr. Markus Osterhoff for help with detector integration and instrumentation, Dr. Michael Sprung for support at the beamline, Jochen Herbst for membrane preparation, and Alessia Kretschmar for help with cell preparation. We acknowledge funding by German Federal Ministry of Education and Research (BMBF) through grant No. 05K19MG2, German Research Foundation (DFG) under Germanys Excellence Strategy -EXC 2067/1-390729940, and Max-Planck School Matter-to-Life.

